# The *Drosophila* Formin Fhod Nucleates Actin Filaments

**DOI:** 10.1101/152348

**Authors:** Aanand A. Patel, Zeynep A. Oztug Durer, Aaron P. van Loon, Margot E. Quinlan

## Abstract

Formins are a conserved group of proteins that nucleate and processively elongate actin filaments. Among them, the formin homology domain-containing protein (FHOD) family of formins contributes to contractility of striated muscle and cell motility in several contexts. However, the mechanisms by which they carry out these functions remain poorly understood. Unlike other formins, mammalian FHOD1 and FHOD3 do not accelerate actin assembly *in vitro*, and have instead been suggested to act as barbed end cappers or bundlers. Here, we show that purified *Drosophila* Fhod, in contrast with the mammalian homologues, potently accelerates actin assembly by nucleation. We found that Fhod binds tightly to barbed ends, where it slows elongation in the absence of profilin and allows elongation in the presence of profilin. Fhod protects barbed ends from capping protein, but dissociates from barbed ends relatively quickly. Finally, we used cosedimentation assays to determine that Fhod binds the sides of actin filaments and bundles filaments. This work establishes that Fhod shares the capacity of other formins to nucleate and bundle actin filaments, but is notably less effective at processively elongating barbed ends.

---

Formins are a major, conserved group of proteins known for their ability to both nucleate new actin filaments and remain processively associated with fast-growing barbed ends via their formin homology 2 (FH2) domains. While bound to the barbed end, many formins accelerate elongation through their proline-rich FH1 domains, which recruit profilin-bound actin monomers to the barbed end. In addition to these classic activities, some formins have additional effects on actin filaments, such as severing, bundling, or cross-linking to microtubules (1). Animals have seven formin families, which share the same domain structure, including the highly conserved FH2 domains. Importantly, formins differ in their actin assembly activities and modes of regulation, allowing them to fulfill distinct cellular roles.

The formin homology domain-containing (FHOD) family of formins has two mammalian isoforms, FHOD1 and FHOD3, which assemble contractile actin structures in several contexts. FHOD1 is widely expressed and assembles stress fibers that contribute to the adhesion, spreading, and motility of numerous cell types (2–8). FHOD3 has also been implicated in the motility of some cancers (9, 10), but its expression is predominantly restricted to striated muscle (11, 12). In cardiomyocytes, FHOD3 localizes to the sarcomere and is required for its assembly and maintenance (12–15). Finally, changes in expression level and polymorphisms of both FHOD1 and FHOD3 are associated with cardiomyopathies (12, 16–19).

The mechanisms by which FHOD family members function remain poorly understood. Unlike other formins, purified FHOD1 and FHOD3 were shown to slow, rather than accelerate, actin assembly in vitro, and have therefore been suggested to act as actin cappers or bundlers (13, 20). In contrast, the cell biology data suggest that FHOD1 and FHOD3 function as actin nucleators in vivo. Across a wide range of cell types, expression of constitutively active FHOD1 or FHOD3 is sufficient to induce the formation of stress fibers (2–5, 13). Furthermore, FHOD3 is required for sarcomere assembly following latrunculin washout (12). These data are most suggestive of actin nucleation, although FHOD1 and FHOD3 might instead function by stabilizing or reorganizing existing actin filaments.

*Drosophila melanogaster* has a single FHOD family member, referred to here as Fhod (also known as Fhos or knittrig). The Fhod gene is alternatively spliced to produce eight different isoforms, which maintain constant FH1 and FH2 domains but alter their regulatory N-termini and C-terminal tails. The role of FHOD proteins in cell motility and contractility is well conserved in *Drosophila*, as Fhod contributes to motility in macrophages and tracheal tip cells (21), sarcomere organization in striated muscle (22, 23), and cardiac contractility (16). Here, we show that purified *Drosophila* Fhod, unlike its mammalian homologues, potently accelerates actin assembly by nucleation. Fhod remains processively associated with the barbed end, where it slows elongation in the absence of profilin and allows elongation, at rates similar to actin alone, in the presence of profilin. Although Fhod does not accelerate barbed end elongation, we find that Fhod protects barbed ends from capping protein with a characteristic run length of about 2 μm. Fhod additionally binds tightly to the sides of filaments and bundles filaments together.

## RESULTS

### Fhod accelerates actin assembly

We purified the C-terminal half of *Drosophila* Fhod isoform A, encompassing the FH1 domain, FH2 domain, and C-terminal tail (Fig 1*A*). This isoform is sufficient to rescue viability in Fhod null flies (21), and its C-terminus is identical to that of isoform H, which rescues sarcomere organization in indirect flight muscle (22). We first tested the effect of Fhod on the assembly of *Acanthamoeba* actin in bulk pyrene assays; Fhod potently accelerated actin assembly in a dose-dependent manner (Fig 1*B-C*). In previous work, purified FHOD1 and FHOD3 did not accelerate assembly of actin from rabbit skeletal muscle (13, 20). Because the activity of some formins depends on the actin isoform (W. T. Silkworth et. al., manuscript submitted), we also tested the ability of Fhod to assemble rabbit muscle actin. We observed similarly accelerated actin assembly with muscle actin and *Acanthamoeba* actin (data not shown). Because Fhod is expressed in both muscle and non-muscle cells, it is logical that Fhod would function equally well with muscle and non-muscle actin. All subsequent experiments were performed with *Acanthamoeba* actin.

**FIGURE 1.**
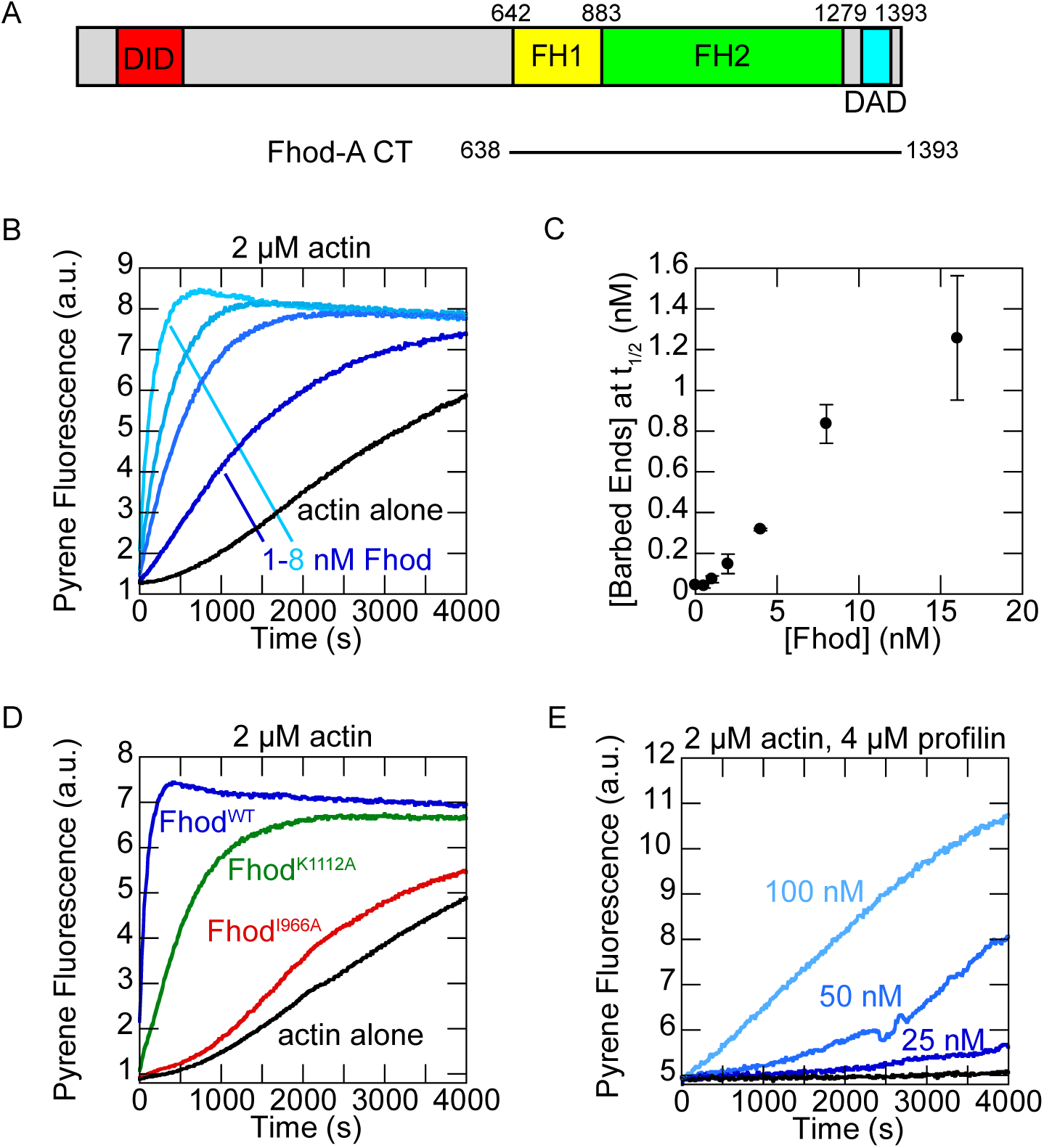
The *Drosophila* formin Fhod accelerates actin assembly. *A*, domain structure of Fhod isoform A. Fhod includes formin homology domains for actin assembly and the Diaphanous inhibitory domain (DID) and Diaphanous autoregulatory domain (DAD) for autoinhibition and regulation. The Fhod construct used in this work spans residues 638-1393 and includes the FH1 domain, FH2 domain, and tail. *B*, assembly of 2 μM *Acanthamoeba castellanii* actin (10% pyrene-labeled) in the absence (black) or presence (blue) of 1-8 nM Fhod. Fhod accelerates actin assembly in a dose-dependent manner. *C*, quantification of barbed end concentrations at the time to half-polymerization from *B*. Data represent the mean ± standard deviation from two independent experiments. *D*, assembly of 2 μM amoeba actin (10% pyrene-labeled) in the presence of 16 nM wild-type (blue) or mutant (I966A, red; K1112A, green) Fhod. Actin assembly by Fhod is reduced moderately by the K1112A mutation and severely by the I966A mutation. *E*, assembly of 2 μM amoeba actin (5% pyrene-labeled) in the presence of 4 μM *S. pombe* profilin and the indicated concentrations of Fhod. Fhod accelerates actin assembly in the presence of profilin.

To further compare Fhod to characterized formins, we introduced two classical mutations, I966A and K1112A, in conserved residues of the FH2 domain (24). The I966A mutation almost completely abolished activity, whereas the K1112A mutation markedly reduced, but did not eliminate, activity (Fig 1*D*). We also tested the ability of Fhod to promote actin assembly in the presence of profilin, which binds most actin monomers in the cell and inhibits spontaneous nucleation. Profilin strongly impaired actin assembly by Fhod, but Fhod still accelerated actin assembly in a dose-dependent manner under these conditions (Fig 1*E*). Fhod is thus similar to other studied formins.

### Fhod promotes actin nucleation but not elongation

Because formins are capable of promoting nucleation as well as elongation, we investigated the relative contributions of these activities. We assessed nucleation activity by imaging actin filaments polymerized in the absence or presence of Fhod. Fhod greatly increased the number of filaments per field of view, indicating that Fhod promotes actin assembly by nucleation (Fig 2*A*-*B*).

**FIGURE 2.**
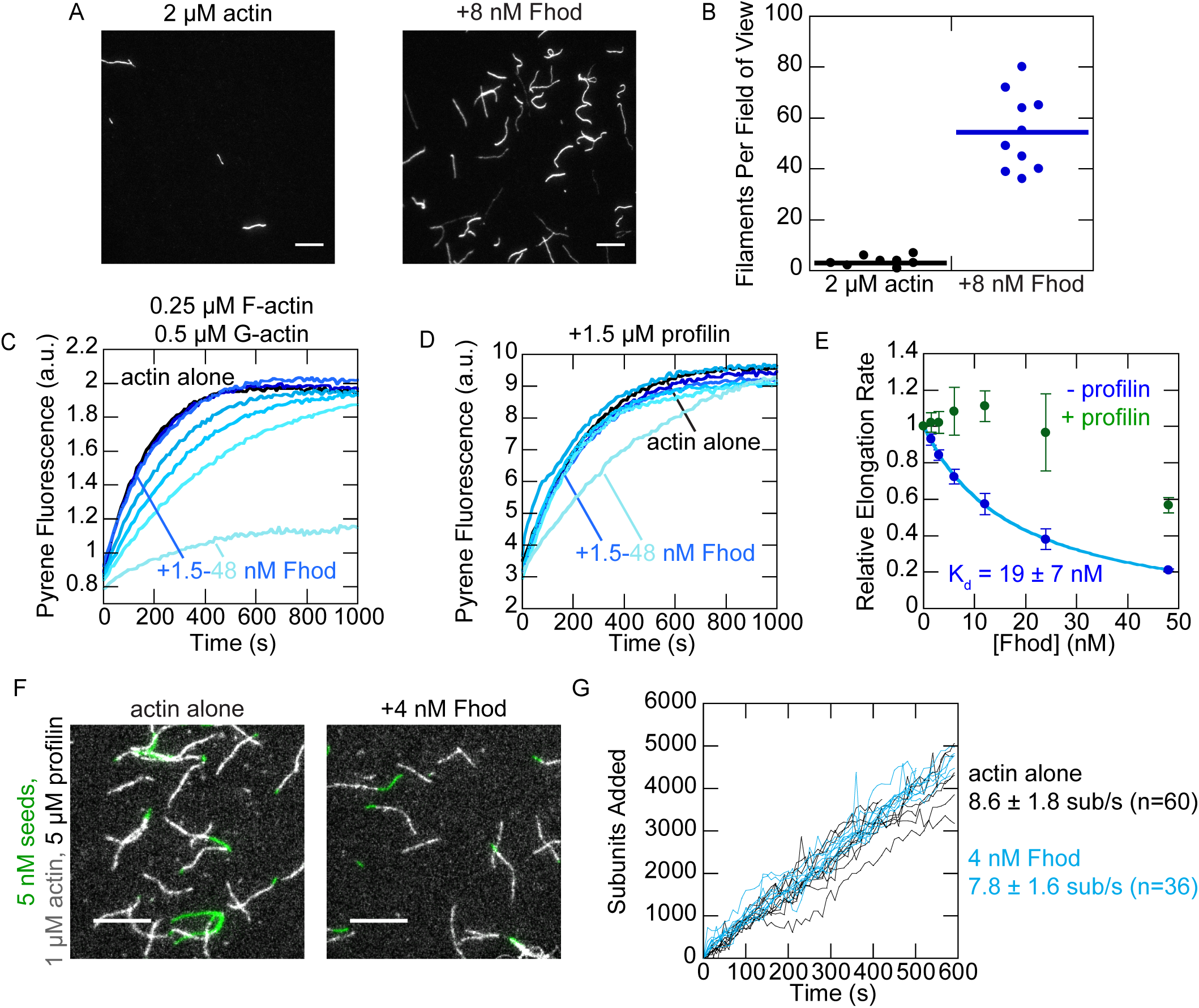
Fhod promotes actin nucleation, but not elongation. *A*, 2 μM actin was polymerized in the absence (left) or presence (right) of 8 nM Fhod for 5 minutes, then stabilized with Alexa Fluor 488- phalloidin and imaged by TIRF microscopy. Scale bar is 10 μm. *B*, quantification of A. Fhod increases the number of actin filaments per field of view. Data are from three independent experiments for each condition. Lines represent the mean. *C*, actin elongation from preformed seeds. Final conditions were 0.25 μM F-actin seeds (~0.4 nM barbed ends), 0.5 μM G-actin (10% pyrene-labeled), in the absence (black) or presence (blue) of 1.5-48 nM Fhod. Fhod slows barbed end elongation in a dose-dependent manner. *D*, actin elongation from preformed seeds, as in C, with 1.5 μM *S. pombe* profilin. Fhod does not slow elongation in the presence of profilin. *E*, quantification of elongation rates from C and D. Elongation rates were measured as the initial slope over the first 90 seconds, relative to the slope of actin alone. Fhod substantially slows elongation only in the absence of profilin. The data and reported K_d_ are the mean ± standard deviation from 3 independent experiments. The binding curve shows the best fit to the average values. *F*, direct observation of barbed end elongation by TIRF microscopy with 5 nM F-actin seeds (1% biotinylated, labeled with Alexa Fluor 647-phalloidin), 1 μM G-actin (10% Alexa Fluor 594-labeled) and 5 μM *S. pombe* profilin, ± 4 nM Fhod. Images were taken 8 minutes after the start of polymerization. Scale bar is 10 μm. *G*, quantification of elongation from *F*. Ten representative traces from each condition are plotted. Elongation rates are average ± standard deviation from 3 flow chambers for each condition.

We then asked how Fhod affects barbed end elongation. Formins typically slow elongation in the absence of profilin and accelerate elongation in the presence of profilin. Using bulk seeded elongation assays, we found that Fhod slows barbed end elongation in the absence of profilin with an affinity of 19 nM (Fig 2*C*, *E*). In the presence of profilin, Fhod had a minimal effect on elongation at most concentrations we tested (Fig 2*D*). At high concentrations of Fhod, the elongation rate appeared to decrease; we attribute this decrease to filament bundling (see below) rather than a true effect on barbed end elongation because the trend does not follow the same dose-dependence we observed in the absence of profilin, and high concentrations of Fhod also caused substantial fluorescence artifacts in actin depolymerization assays. To confirm that Fhod does not alter barbed end elongation in the presence of profilin, we directly measured elongation rates using total internal reflection fluorescence (TIRF) microscopy. In agreement with our bulk assays, we detected no difference in elongation rates (Fig 2*F*-*G*).

Because we did not observe clear evidence of processive elongation by Fhod, we used several additional assays to verify and characterize the interaction between Fhod and barbed ends. We first verified barbed end binding in bulk barbed end depolymerization assays. The dose dependence gives us an additional measure of the affinity between Fhod and barbed ends. Fhod inhibited actin depolymerization with a K_d_ of 5 nM, similar to our measurement from the seeded elongation assays (Fig 3*A-B*). We then used actin reannealing assays, in which two colors of sheared actin filaments were mixed and allowed to reanneal. Fhod inhibited actin reannealing, indicating that it binds barbed ends and can remain bound on the timescale of minutes (Fig 3*C*). Thus Fhod binds barbed ends tightly, like other formins, but does not accelerate elongation, unlike other formins.

**FIGURE 3.**
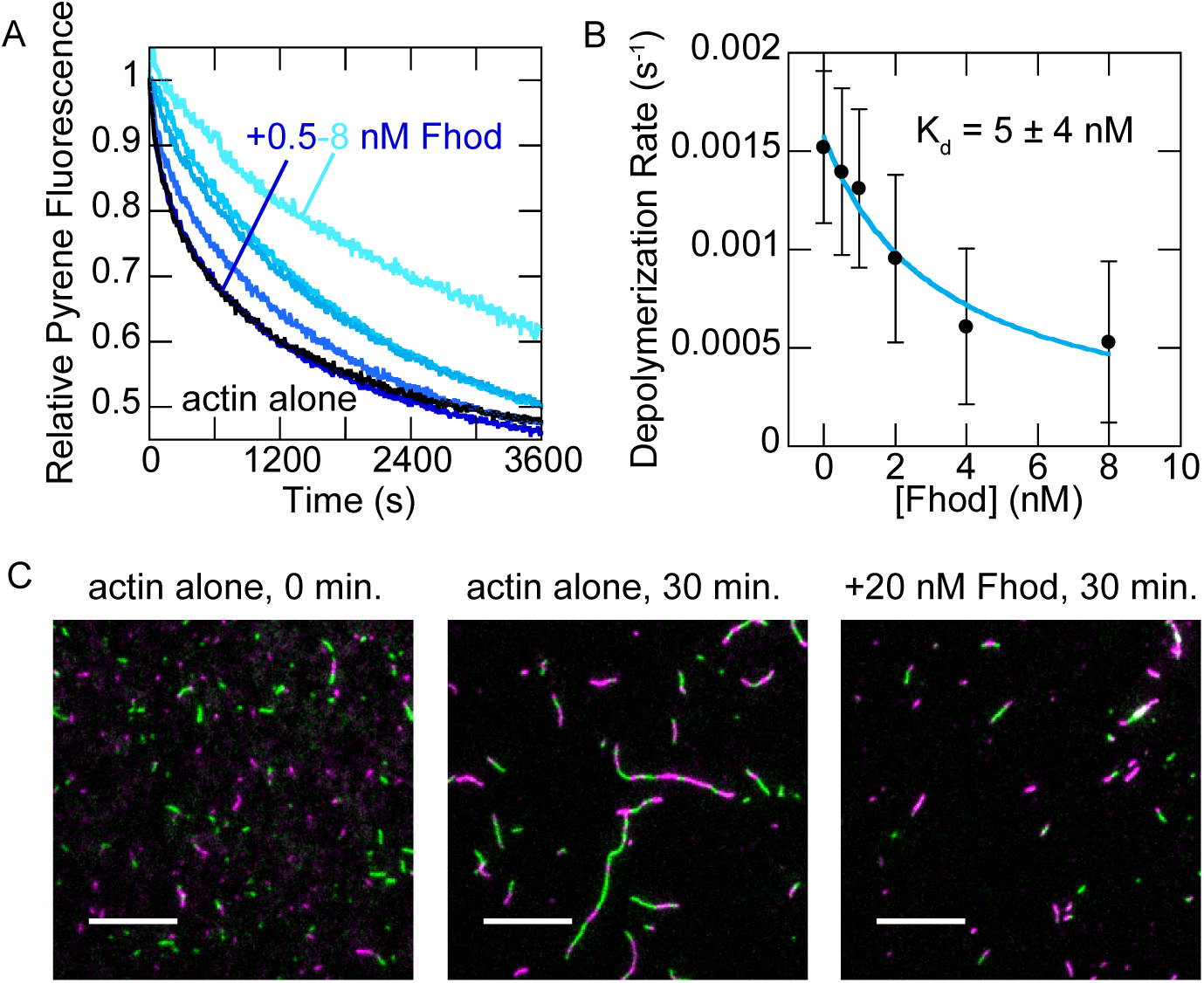
Fhod binds barbed ends. *A*, actin filaments were depolymerized by dilution to 0.1 μM (70% pyrene-labeled) with varying concentrations of Fhod. Data were normalized against the initial pyrene fluorescence. *B*, quantification of depolymerization rates from *A*, represented by the initial slope over the first 90 seconds. Fhod binds barbed ends and slows depolymerization. The data and reported K_d_ are the mean ± standard deviation from six independent experiments. The binding curve shows the best fit to the average values. *C*, Actin filaments, stabilized with Alexa Fluor 488- (green) or rhodamine-(magenta) phalloidin, were sheared and allowed to reanneal for 30 minutes. Final concentrations were 0.25 μM F-actin ± 10 nM Fhod. Fhod inhibits reannealing of the sheared filaments. Scale bar is 10 μm.

### Fhod antagonizes capping protein

We measured the ability of Fhod to antagonize capping protein, which binds tightly to barbed ends and prevents elongation. In bulk seeded elongation assays, 6 nM capping protein was sufficient to completely abolish actin elongation. Fhod abrogated this effect when added to F-actin seeds prior to capping protein (Fig 4*A*). We fit these data to a competition binding equation to determine that Fhod has an apparent K_d_ of 7 nM for growing barbed ends (Fig 4*B*), consistent with our previous measurements. Recent evidence suggests that formins can antagonize capping protein not only by passive competition for the barbed end, but also by binding capped barbed ends and actively displacing capping protein (25, 26). However, filaments did not grow when capping protein was added before Fhod (data not shown), indicating that the actin elongation we observed was due to Fhod processively protecting the barbed end, and not due to Fhod actively displacing capping protein from barbed ends.

**FIGURE 4.**
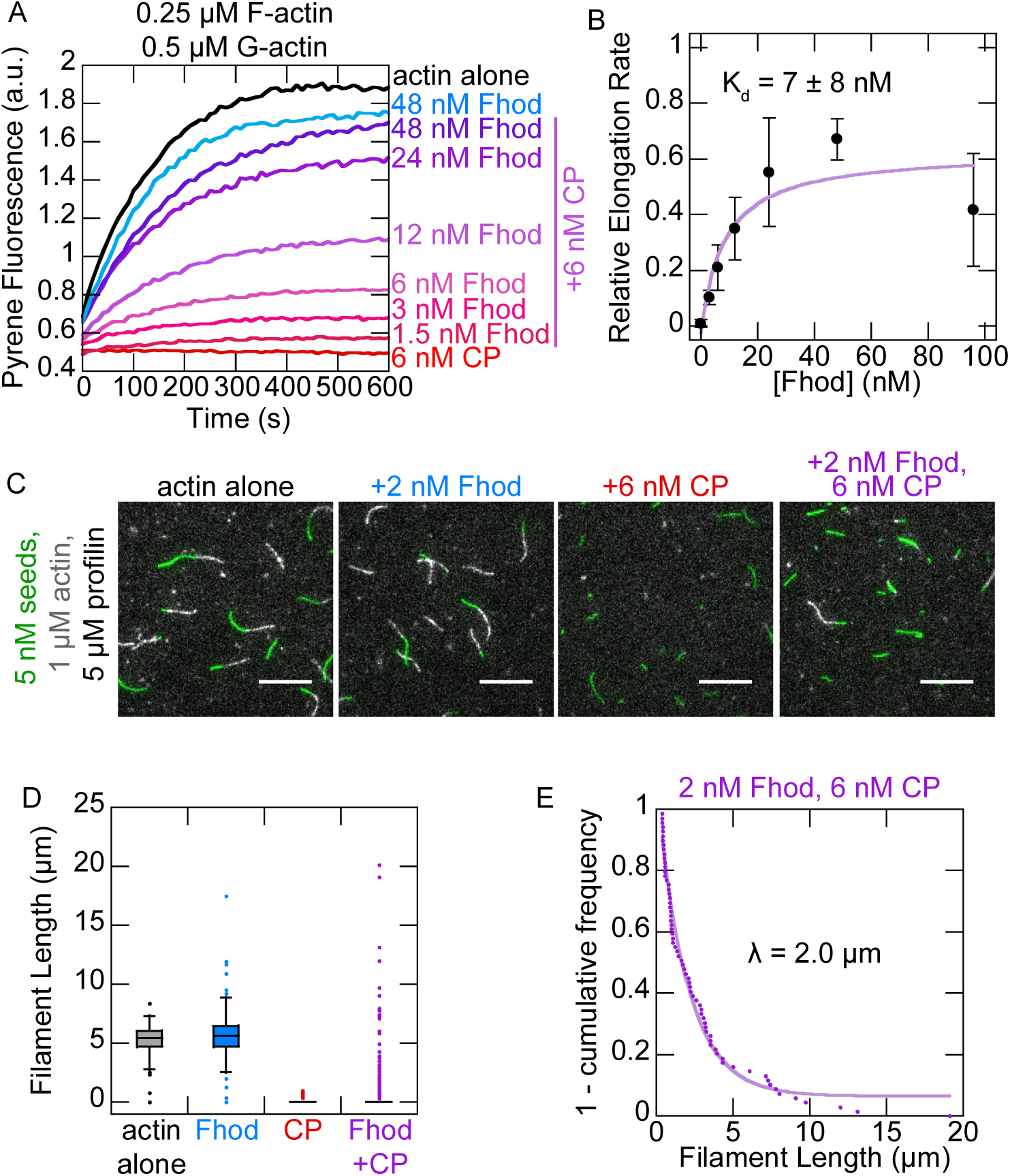
Fhod antagonizes capping protein. *A*, actin elongation from preformed seeds with a range of Fhod concentrations added before capping protein. Final conditions were 0.25 μM F-actin seeds (~0.4 nM barbed ends), 0.5 μM G-actin (10% pyrene-labeled), 1.5 μM *S. pombe* profilin, ± 6 nM mouse capping protein and 1.5-48 nM Fhod. *B*, quantification of elongation rates from *A*, measured as the initial slope over the first 90 seconds relative to the slope of actin alone. Fhod antagonizes capping protein, allowing elongation. Data with Fhod and capping protein were fit to a competition binding model to determine the affinity of Fhod to barbed ends. The data and reported K_d_ are mean ± standard deviation from four independent experiments. The binding curve shows the best fit to the average values. *C*, observation of actin elongation (white) from preformed seeds (green) with Fhod added before capping protein. Final conditions were 5 nM F-actin seeds (1% biotinylated, labeled with Alexa Fluor 647-phalloidin), 1 μM G-actin (10% Alexa Fluor 594-labeled), 5 μM *S. pombe* profilin, ± 2 nM Fhod and 6 nM capping protein. Images were taken five minutes after the start of polymerization. Scale bar is 10 μm. *D*, quantification of filament lengths from *C*. Data represent the amount of elongation from preformed seeds (n > 150 for each condition). At least 5 fields of view from 1 (actin alone) or 2 (all other conditions) flow chambers were analyzed for each condition. In conditions with capping protein, no box is visible because over 75% of seeds did not elongate. *E*, exponential fit of filament lengths in the presence of both Fhod and capping protein from *D*, excluding seeds that did not elongate (n=69 filaments from 2 flow chambers).

We used TIRF microscopy to observe competition between Fhod and capping protein on individual filaments. We incubated seeds with Fhod prior to adding capping protein and actin monomers, then measured how long Fhod could protect the growing barbed end. Consistent with our bulk assays, we found that barbed ends were completely capped by capping protein in the absence of Fhod, but were able to grow in the presence of Fhod. Because the vast majority of filaments were capped by the time we could start imaging (2-3 minutes after the start of polymerization), we measured filament lengths in static images taken 5 minutes after addition of actin monomers (Fig 4*C*-*D*). By fitting the filament lengths to a single exponential curve, we determined that Fhod has a characteristic run length of 2 μm (Fig 4*E*). This provides us with an approximate measure of Fhod’s processivity, with two assumptions: (1) capping protein binds to barbed ends as soon as Fhod dissociates, and (2) capping protein does not cap barbed ends that are already bound by Fhod. The first assumption is consistent with the strong affinity (~0.4 nM) of capping protein for barbed ends. However, the ability of capping protein to bind mDia1-bound barbed ends and displace mDia1 (25, 26) suggests that capping protein might also bind Fhod-bound barbed ends, which would make our measurement of Fhod’s processivity an underestimate.

### Fhod bundles actin filaments

Finally, we asked whether Fhod shares the capacity of other formins to bind the sides of actin filaments and bundle them. In high speed cosedimentation experiments, Fhod pelleted with F-actin, demonstrating that Fhod binds the sides of actin filaments, with a K_d_ of 180 nM (Fig 5*A*). In low-speed cosedimentation, Fhod increased the amount of F-actin in the pellet, indicating that it forms actin bundles (Fig 5*B*). We used TIRF microscopy to determine the polarity of actin bundles formed by Fhod. Actin bundles typically elongated from both ends, indicating that the actin filaments in these bundles were not arranged exclusively in parallel (Fig 5*C*).

**FIGURE 5.**
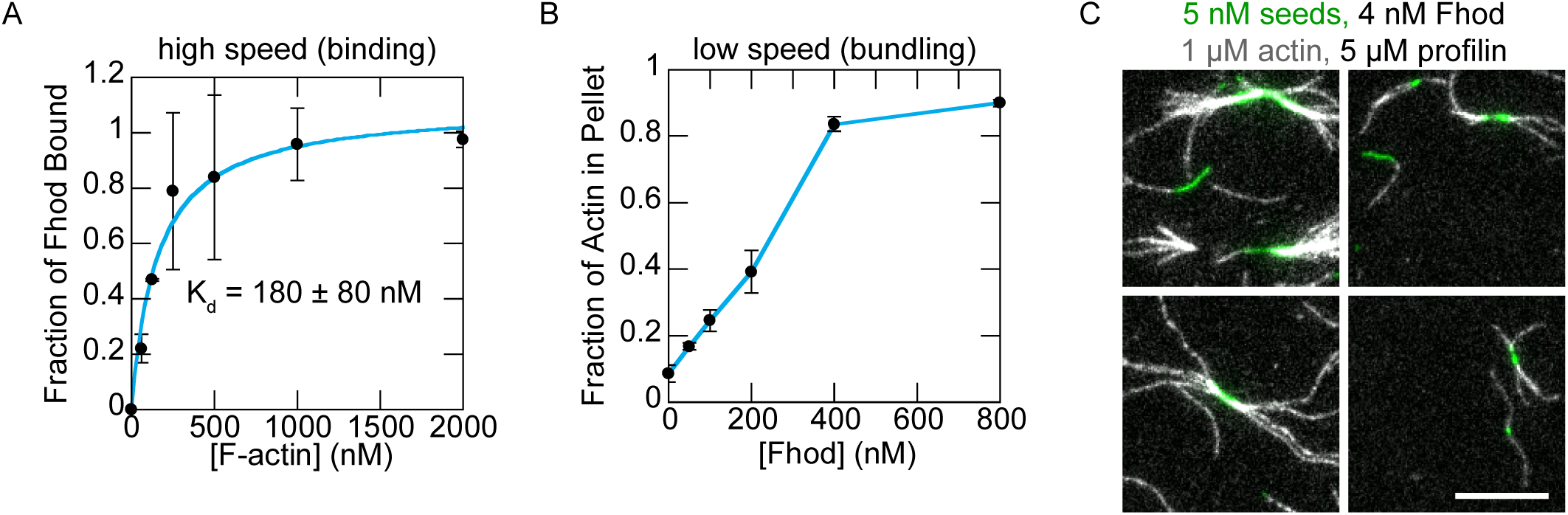
Fhod binds the sides of actin filaments and forms actin bundles. *A*, high speed cosedimentation assay with 250 nM Fhod and varying concentrations of phalloidin-actin. The fraction of Fhod bound to F-actin was quantified by measuring the amount of Fhod in the pellet and correcting for the amount of Fhod that pellets in the absence of F-actin. The data and reported K_d_ are mean ± standard deviation from three independent experiments. The binding curve shows the best fit to the average values. *B*, low speed cosedimentation assay with 5 μM phalloidin-actin and varying concentrations of Fhod. Data are mean ± standard deviation from two independent experiments. *C*, representative images of Fhod-induced actin bundles visualized by TIRF microscopy. Final conditions were 5 nM F-actin seeds (1% biotinylated, labeled with Alexa Fluor 647-phalloidin), 1 μM G-actin (10% Alexa Fluor 594-labeled), 5 μM *S. pombe* profilin, 4 nM Fhod. Bundles elongate at both ends, indicating mixed or antiparallel polarity. Scale bar is 10 μm.

## DISCUSSION

Here, we show that *Drosophila* Fhod shares the classic activities of formins, actin nucleation and processive elongation, with the additional capacity to bundle actin filaments. These observations differ substantially from those reported for mammalian FHOD1 and FHOD3. Whereas both FHOD1 and FHOD3 were reported to slow actin assembly (13, 20), we found that *Drosophila* Fhod is a potent actin nucleator similar to other formins. Our ability to achieve high expression in *E. coli*, in contrast with the requirement of a eukaryotic expression system for human FHOD1 (20), indicates a difference in the folding, conformation, or stability of the two proteins. Therefore, we expect that FHOD1 and FHOD3 might function as actin nucleators *in vivo*, despite their inability to accelerate actin assembly *in vitro*. FHOD1 and FHOD3 induce the formation of stress fibers in several cell types (2–5, 13), and FHOD3 is required for sarcomere assembly (12). These functions are well conserved in *Drosophila* (21, 22) and most consistent with actin nucleation. However, it remains formally possible that FHOD proteins instead stabilize or bundle filaments that are polymerized by a different actin nucleator. Future work will be needed to determine what accounts for the differences we observe between *Drosophila* and mammalian FHOD proteins.

We did not observe evidence of accelerated barbed end elongation with Fhod. This is not unprecedented, as some formins such as *Drosophila* Daam (27) and mouse FMNL1 (28) either slow barbed end elongation or leave the elongation rate unchanged in the presence of profilin. Mammalian FHOD1 and FHOD3 were also shown to slow barbed end elongation, but in a profilin-independent manner (13, 20). We expect that both the FH1 and FH2 domains contribute to Fhod’s inability to accelerate barbed end elongation. The slow barbed end elongation in the absence of profilin is suggestive of an FH2 domain that spends most of the time in a closed conformation, similar to Cdc12 (29). The addition of profilin restores the elongation rate to that of actin alone, indicating that the FH1 domain has some ability to recruit profilin-actin, but perhaps not as effectively as the FH1 domains of other formins. The effectiveness of polyproline tracks in the FH1 domain depend on the number of prolines and their distance from the FH2 domain (30, 31). The polyproline tracks of the Fhod FH1 domain are located relatively far from the FH2 domain, with the closest track (PPPMMPP) located 31 residues from the FH2 domain. For comparison, the weak elongator Cdc12 has its closest polyproline track 26 residues away from the FH2 domain, whereas the strong elongators Bni1 and mDia1 have their closest polyproline tracks 22 and 16 residues away, respectively.

We approximate that Fhod has a characteristic run length of 2 μm, which is equivalent to a dissociation rate of ~0.01 s^−1^ based on the elongation rate of 8 subunits/second. This dissociation rate is an order of magnitude faster than mDia1 and several orders of magnitude faster than mDia2, Bni1, Cdc12, and Capu (32, 33), which does not fit the general trend of faster elongation rates resulting in faster dissociation rates (30). We observed evidence of Fhod protecting barbed ends only when Fhod was incubated with seeds and actin monomers in a tube prior to introducing the mixture onto the surface. This suggests that the surface hinders Fhod’s processivity, making our measurement of Fhod’s processive elongation activity an underestimate. However, that Fhod is sensitive to conditions that do not perturb the processivity of other formins may indicate that processive elongation is not a critical activity of Fhod. FHOD family members generally localize to the relatively short actin filaments found in stress fibers and the sarcomere, which likely do not require accelerated barbed end growth. Therefore, Fhod’s nucleation and bundling activities might be more important in these contexts. We found that Fhod is indeed a potent actin bundler; its affinity of 0.18 μM for sides of actin filaments is comparable to the formins Fus 1(34) and AFH1 (35), and an order of magnitude stronger than mDia1 (36), Daam (27), and Capu (33). Alternatively, it is possible that Fhod accelerates actin elongation *in vivo* through collaborations with other proteins; for example, CLIP-170 was recently shown to augment the processive elongation of mDia1 (37).

## EXPERIMENTAL PROCEDURES

### Protein expression, purification, and labeling

cDNA for *Drosophila* Fhod isoform A (SD08909, obtained from FlyBase) was used as a template to clone the C-terminal half of Fhod (residues 638-1393) into a modified version of the pET15b plasmid with an N-terminal 6xHis tag. Point mutations were generated by site-directed mutagenesis as described (38). Fhod constructs were transformed in Rosetta(DE3) cells (Novagen), which were grown in 1 liter of Terrific Broth supplemented with 100 mg/L ampicillin and 32 mg/L chloramphenicol. When the OD reached ~0.8, expression was induced by adding 0.25 mM IPTG and shaking overnight at 18 °C. Cells were harvested by centrifugation, washed in PBS, and flash frozen in liquid nitrogen.

Cell pellets were thawed in extraction buffer (10 mM MOPS pH 7, 0.5% Triton X-100, 1 mM DTT, 1 mM PMSF, 2 μg/mL DNaseI). All subsequent steps were performed on ice or at 4 °C. Cells were lysed by microfluidizing, cleared by centrifugation at 20,000 x *g* for 20 minutes, then purified using a HitrapSP-FF cation exchange column (GE Life Sciences) with a gradient of 0.3-1 M NaCl over 16 column volumes. Pooled fractions were diluted at least six-fold into 10 mM Tris pH 8, 1 mM DTT and further purified on a MonoQ anion exchange column (GE Life Sciences) with a gradient of 40-500 mM NaCl over 50 column volumes. Peak fractions were exchanged into storage buffer (10 mM Tris pH 8, 150 mM NaCl, 1 mM DTT, 0-20% glycerol), centrifuged at 20,000 x *g* for 20 minutes, flash frozen in liquid nitrogen, and stored at −80 °C. Fhod protein concentrations were determined using the absorbance at 280 nm with an extinction coefficient of 122,840 M^−1^ cm^−1^ (ProtParam), which was verified by comparing the absorbances of native and denatured protein. All Fhod concentrations are reported in terms of dimer concentrations.

*Acanthamoeba castellani* actin was purified (39) and labeled with Alexa Fluor 594 succinimidyl ester (40), pyrene iodoacetamide (39), or EZ-link maleimide-PEG2-biotin (Thermo Scientific) (41) according to published protocols. *S. pombe* profilin was purified as described (28) on a polyproline column kindly provided by the Reisler lab (University of California, Los Angeles). The concentration was determined using the absorbance at 280 nm with an extinction coefficient of 1.63 OD/mg/mL (42). Mouse capping protein was purified as described (43).

### Pyrene assays

Pyrene assays were performed essentially as described (44) on an Infinite 200 Pro plate reader (Tecan). In all assays, Fhod was diluted in storage buffer before addition to polymerization buffer to improve stability. The concentration of barbed ends was calculated from the slope (obtained by linear regression over 90 seconds) using the equation [be] = elongation rate/(k_+_[G-actin] - k_-_), where k_+_ = 11.6 μM^−1^ s^−1^ and k_-_ = 1.4 s^−1^ (45). For seeded elongation assays, actin filaments were sheared by passing three times through a 24-gauge needle, then aliquoted into each well of a clear microplate. Proteins were added to the seeds and incubated for 2-4 minutes at room temperature; in experiments with both Fhod and capping protein, capping protein was added 2-4 minutes after addition of Fhod. Seeds and additional proteins in KMEH (10 mM HEPES pH 7, 1 mM EGTA, 50 mM KCl, 1 mM MgCl_2_) were added to Mg-actin to initiate actin elongation. Elongation rates were determined by linear regression over the first 90 seconds and normalized against the rate of actin alone in each experiment. For experiments without capping protein, the affinity of Fhod for barbed ends was determined by fitting the data to the simplified binding equation, r = [Fhod] / ([Fhod] + K_d_) * a + b, where r is the normalized elongation rate. Data for capping protein experiments were analyzed using a competition binding model (46). The following equation was used to determine the K_d_ of Fhod for growing barbed ends:

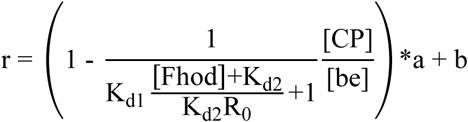

where r is the normalized elongation rate, K_d1_ is the affinity of capping protein for barbed ends (0.4 nM, measured in a seeded elongation assay in the absence of Fhod), K_d2_ is the affinity of Fhod for barbed ends, and R_0_ is the concentration of free barbed ends when [Fhod]=0. The total concentration of barbed ends was calculated from the initial slope of the polymerization trace for actin alone as described above.

For depolymerization assays, 1 μM F-actin (70% pyrene-labeled) was incubated for at least 15 minutes at room temperature, then depolymerized by diluting 10-fold in 1x KMEH with additional proteins. The depolymerization rate was determined by linear regression over the first 90 seconds. The affinity of Fhod for barbed ends was determined by fitting the data to the simplified binding equation as above.

### TIRF microscopy

In nucleation assays, assembly of 2 μM actin was initiated by the addition of KMEH with or without Fhod. After five minutes, actin was removed from the reaction and stabilized by diluting 10-fold in 1x KMEH supplemented with Alexa Fluor 488-phalloidin. Actin was incubated with phalloidin for ten minutes, diluted 20-fold in 1x KMEH supplemented with 100 mM DTT, spotted on a poly-L-lysine-coated coverslip, and imaged. All steps were performed as delicately as possible with a cut pipet tip to minimize shearing.

For elongation experiments, biotinylated coverslips were prepared as follows. Coverslips were rinsed three times in MilliQ water, placed in 2% Hellmanex (Hellma Analytics) at 60-65 °C for 2 hours, then rinsed another five times in MilliQ water. Once dry, the coverslips were silanized with GOPTS for 1 hour in a hybridization oven. Unreacted GOPTS was removed by rinsing three times with acetone. Coverslips were then PEGylated with a mixture of methoxy-PEG-NHS and biotin-PEG-NHS as described (44).

Flow chambers of about 15 μL were assembled on the slide using strips of double-sided tape. Flow chambers were prepared with the following steps: 1) block with 25 μL 1% Pluronic F-127 (Sigma), 50 μg/mL casein, in PBS, for 2 minutes; 2) 25 μL 1x KMEH; 3) 25 μL 40 nM streptavidin in 1x KMEH; 4) 25 μL 1x TIRF buffer [1x KMEH, 0.5% methylcellulose (400 cP, Sigma), 50 mM DTT, 0.2 mM ATP, 20 mM glucose]; 5) 50 μL Mg-actin and additional proteins to be assayed, in 1x TIRF buffer supplemented with 5 nM F-actin seeds (1% biotinylated, stabilized with Alexa Fluor 647- phalloidin), 250 μg/mL glucose oxidase, 50 μg/mL catalase, and 50 μg/mL casein. Fhod was incubated with seeds for at least 30 seconds prior to addition of Mg-actin; in experiments with both Fhod and capping protein, Fhod was incubated with seeds for 15 seconds prior to addition of capping protein, and Mg-actin was added after an additional 30 seconds.

To determine the characteristic run length of Fhod on barbed ends in the presence of capping protein, 1 - cumulative frequency was treated as the fraction of filaments that were still elongating at a particular length. The data were fit to the exponential equation, 1 - cf = e^−l/λ^ * a + b, where cf is the cumulative frequency, l is the filament length, and λ is the characteristic run length.

Reannealing assays were conducted essentially as described (47) using Alexa Fluor 488- or rhodamine-labeled phalloidin-actin, sheared by passing three times through a 27-gauge needle. Final concentrations were 250 nM F-actin and 10 nM Fhod. Samples were diluted 50-fold, spotted on poly-L-lysine-coated coverslips, and imaged.

In all experiments, actin filaments were visualized on a DMI6000 TIRF microscope (Leica) with an HCX PL APO objective (100x magnification, N.A.=1.47), and an Andor DU-897 camera, using the Leica Application Suite Advanced Fluorescence software. Experiments were conducted at room temperature. Images were obtained at 10 second intervals for 10 minutes. Images were processed by applying rolling ball background subtraction and a Gaussian filter. Filament lengths were quantified using the JFilament plugin for Fiji (48).

### Cosedimentation

For high speed cosedimentation, 250 nM Fhod was incubated with varying concentrations of phalloidin-stabilized F-actin for 30 minutes at room temperature. Samples were centrifuged at 90,000 rpm for 25 minutes in a TLA-100 rotor. Pellets were concentrated by resuspending in one fourth the original volume. Supernatants and pellets were analyzed with SDS-PAGE, and gels were stained with SyproRed. The amount of Fhod in each fraction was quantified using QuantityOne software, dividing the intensities of pellet bands by four to correct for the four-fold concentration of pellets with resuspension. The fraction of Fhod bound to F-actin was calculated by adjusting for the amount of Fhod that pellets in the absence of F-actin with the following equation: θ = (p − p_0_) / (1 − p_0_), where θ is the fraction of Fhod bound to F-actin, p is the fraction of Fhod in the pellet, and p0 is the fraction of Fhod in the pellet in the absence of F-actin. The affinity of Fhod for F-actin was determined by fitting the data to the binding equation, θ = [F-actin] / ([F-actin] + K_d_) * a + b.

For low speed cosedimentation, Fhod was incubated with phalloidin-stabilized actin filaments (final concentration of 5 μM) for 1 hour at room temperature, then centrifuged at 12,000 x g for 15 minutes. The amount of actin in the supernatants and pellets was quantified by Coomassie-staining SDS-PAGE gels.

## Acknowledgments

We thank members of the Quinlan and Reisler labs for discussions and feedback.

## Conflict of interest

The authors declare that they have no conflicts of interest with the contents of this article.

## Author contributions

All authors contributed to design of the experiments. A.A.P., Z.A.O.D., and A.P.v.L. performed experiments. A.A.P. and M.E.Q. wrote the manuscript with revisions from Z.A.O.D. and A.P.v.L.

## FOOTNOTES

This work was supported by the following grants from the National Institutes of Health: NRSA F30 HL137263 (A.A.P.), UCLA Medical Scientist Training Program T32 GM008042 (A.A.P.), and R01 GM096133 (M.E.Q.). The content is solely the responsibility of the authors and does not necessarily represent the official views of the National Institutes of Health.

The abbreviations used are: FHOD, formin homology domain-containing protein; TIRF, total internal reflection fluorescence; FH, formin homology domain; IPTG, isopropyl β-D-thiogalactoside; DTT, dithiothreitol; PMSF, phenylmethanesulfonyl fluoride; EGTA, ethylene glycol tetraacetic acid; GOPTS, (3-glycidyloxypropyl)trimethoxysilane

